# The dual role of the RETINOBLASTOMA-RELATED protein in the DNA damage response is spatio-temporally coordinated by the interaction with LXCXE-containing proteins

**DOI:** 10.1101/2021.12.22.473497

**Authors:** Jorge Zamora Zaragoza, Katinka Klap, Renze Heidstra, Wenkun Zhou, Ben Scheres

## Abstract

Living organisms face threats to genome integrity caused by environmental challenges or metabolic errors in proliferating cells. To avoid the spread of mutations, cell division is temporarily arrested while repair mechanisms deal with DNA lesions. Afterwards, cells either resume division or respond to unsuccessful repair by withdrawing from the cell cycle and undergoing cell differentiation or cell death. How the success rate of DNA repair connects to the execution of cell death remains incompletely known, particularly in plants. Here we provide evidence that the Arabidopsis thaliana RETINOBLASTOMA-RELATED1 (RBR) protein, shown to play structural and transcriptional functions in the DNA damage response (DDR), coordinates these processes in time by successive interactions through its B-pocket sub-domain. Upon DNA damage induction, RBR forms nuclear foci; but the N849F substitution in the B-pocket, which specifically disrupts binding to LXCXE motif-containing proteins, abolishes RBR focus formation and leads to growth arrest. After RBR focus formation, the stress-responsive gene *NAC044* arrests cell division. As RBR is released from nuclear foci, it can be bound by the conserved LXCXE motif in NAC044. RBR-mediated cell survival is inhibited by the interaction with NAC044. Disruption of NAC044-RBR interaction impairs the cell death response but is less important for NAC044 mediated growth arrest. Noteworthy, unlike many RBR interactors, NAC044 binds to RBR independent of RBR phosphorylation. Our findings indicate that the availability of the RBR B-pocket to interact with LXCXE-containing proteins couples RBR DNA repair functions and RBR transcriptional functions of in the cell death program.

## Introduction

Living organisms encounter daily challenges to genome integrity that jeopardize survival and reproduction. In response to intrinsic or environmental factors (Chen et al., 2019; Tsegay et al., 2019; Yi et al., 2014), eukaryotic DNA damage triggers a variety of responses collectively known as DNA damage response (DDR). The ATM/ATR kinases initiate a phosphorelay system to tag the damage site, pause the cell cycle, and repair the lesion. If the damage is too extensive to repair, cells activate a suicidal program –apoptosis in animals, and Programmed Cell Death (PCD) in plants – to avoid propagation of mutations (Chen, 2016; Ciccia and Elledge, 2010; Hu et al., 2016; Kim et al., 2019; Lanz et al., 2019; Waterworth et al., 2019). In animals, DDR is largely mediated by p53, a transcription factor that activates the DNA repair machinery and the transcriptional responses leading to cell division arrest and, if necessary, senescence and apoptosis (Chen, 2016; Kastenhuber and Lowe, 2017; Williams and Schumacher, 2016). In plants, no p53 orthologs have been found, and DDR relies on the functional analog SUPPRESSOR OF GAMMA RESPONSE1 (SOG1), a member of the plant-specific NAC-transcription factor family (NAM-ATAF-CUC (Bourbousse et al., 2018; Mahapatra and Roy, 2020; Yoshiyama et al., 2009).

Previous studies identified direct and indirect targets of SOG1 upon DNA damage in Arabidopsis thaliana (Bourbousse et al., 2018; Ogita et al., 2018). Besides activating the majority of the DNA repair genes, SOG1 represses cell cycle and induces cell death by directly activating NAC044 and NAC085, the closest SOG1 paralogs (Bourbousse et al., 2018; Ogita et al., 2018; Takahashi et al., 2019). NAC044/NAC085 stabilize MYB3R3 repressor proteins that in turn bind to the MSA sequence present in G2/M gene promoters, arresting cell division (Chen et al., 2017; Kobayashi et al., 2015; Takahashi et al., 2019). However, how NAC044/NAC085 induce cell death in after DNA injury is unclear.

In recent years, another cell cycle regulator emerged as a central player in the DDR acting in parallel to SOG1: the transcriptional repressor RETINOBLASTOMA-RELATED1 protein (RBR), which is the homolog of the human tumor suppressor pRB and a major regulator of the G1/S phase transition (Biedermann et al., 2017; Bouyer et al., 2018; Cruz-Ramírez et al., 2012; Horvath et al., 2017). RBR is a multifunctional protein that integrates environmental information into cell cycle and developmental programs by interacting with a plethora of transcriptional and chromatin regulators (Cruz-Ramírez et al., 2012; Desvoyes et al., 2014a; Gutierrez, 2005; Gutzat et al., 2012; Harashima and Sugimoto, 2016; Johnston et al., 2008; Matos et al., 2014)(Borghi et al., 2010; Chen et al., 2011; Cruz-Ramírez et al., 2012; Harashima and Sugimoto, 2016; Perilli et al., 2013). In proliferating cells, RBR binds to E2F-DP heterodimeric transcription factors to prevent S-phase onset until Cyclin D-CDKA kinases phosphorylate RBR to release E2F-DP, allowing cell cycle progression (Berckmans and De Veylder, 2009; Desvoyes et al., 2014b; Magyar et al., 2012; Polit et al., 2012; De Veylder et al., 2002).

Upon DNA damage, RBR plays both a transcriptional and a structural role. It regulates the expression of repair genes and mediates the localization of the RAD51 repair protein to DNA damage sites, visualized as nuclear foci where other proteins such as E2FA and BRCA1 colocalize with RBR (Biedermann et al., 2017; Bouyer et al., 2018; Horvath et al., 2017). Reduction of RBR levels leads to genome instability, cell death, and hypersensitivity to DNA damaging treatments (Biedermann et al., 2017; Cruz-Ramírez et al., 2013; Horvath et al., 2017). Thus, it is likely that RBR participates in the mechanism mediating PCD following DNA damage.

RBR belongs to the ‘pocket protein’ family, characterized by the A- and B-pocket subdomains that fold into the ‘pocket domain’, a N-domain, a C-terminal region, and multiple sites for CDK-mediated phosphorylation (Desvoyes and Gutierrez, 2020; Dick and Rubin, 2013; Gutzat et al., 2012; Rubin, 2013). Pocket-protein functions rely on their ability to form protein interactions regulated by phosphorylation (Dick and Rubin, 2013; Narasimha et al., 2014; Sanidas et al., 2019). E2Fs bind to the A-B subdomains interface, while Cyclin D proteins and many others bearing the conserved LXCXE motif dock in the LXCXE-binding cleft located at the B-subdomain (DeCaprio, 2009; Flemington et al., 1993; Helin et al., 1993). A point mutation that specifically disrupts pRB-LXCXE interactions fails to irreversibly arrest cell division in human cell lines (Chen and Wang, 2000), and hampers anti-tumorigenic activity of pRB after induced DNA damage in mice (Bourgo et al., 2011). Here, we used the same amino acid change to dig deeper into the molecular determinants underlying Arabidopsis thaliana RBR roles during DDR. We show that the ability of RBR to interact with LXCXE-containing proteins is crucial to withstand DNA damage. An as yet unidentified factor recruits RBR to nuclear foci early after DNA damage induction. Following clearance from foci, RBR interacts with NAC044 in a LXCXE dependent and phosphorylation-independent manner. Specific disruption of RBR-NAC044 interaction revealed that the SOG1 and RBR pathways converge on the transcriptional regulation of cell death. Collectively, our results support the existence of an LXCXE-coordinated timeframe for the dual RBR role during the DDR.

## Results

### RBR interacts with LXCXE proteins to function in the DNA damage response

The ability of mouse pRB to interact with LXCXE proteins is dispensable under ideal growth conditions, but it is essential when DNA is damaged (Bourgo et al., 2011). To investigate the role of RBR-LXCXE interactions in the plant DDR, we analysed the effect of mutating asparagine (N) 849 to phenylalanine (F) in Arabidopsis RBR, hereafter referred to as RBR^NF^ (Cruz-Ramirez et al., 2013). N849 is a conserved residue of the RBR LXCXE-binding cleft located in the B-pocket subdomain (Gutzat et al., 2012) (Fig S1), and other studies in mammals (Bourgo et al., 2011; Chen and Wang, 2000) and Arabidopsis (Cruz-Ramírez et al., 2013) have used the NF allele to disrupt LXCXE interactions. We transformed pRBR::RBR^NF^:vYFP (RBR^NF^-YFP) into our previously reported amiGO-RBR line (hereafter amiGO), an RBR-targeted amiRNA that permits complementation with transgenic RBR lacking the 3’UTR (Cruz-Ramírez et al., 2013).

While the roots of both variants grew normally in standard conditions (Fig 1A), only RBR-YFP was able to sustain growth on medium supplemented with zeocin (Fig 1B, Fig 1SB), a radiomimetic genotoxic agent that creates double strand breaks (DSB) in the DNA (Kim et al., 2019). A recovery period after zeocin treatment revealed that the RBR^NF^ allele was unable to cope with with DNA damage (Fig 1C).

**Figure 1.**
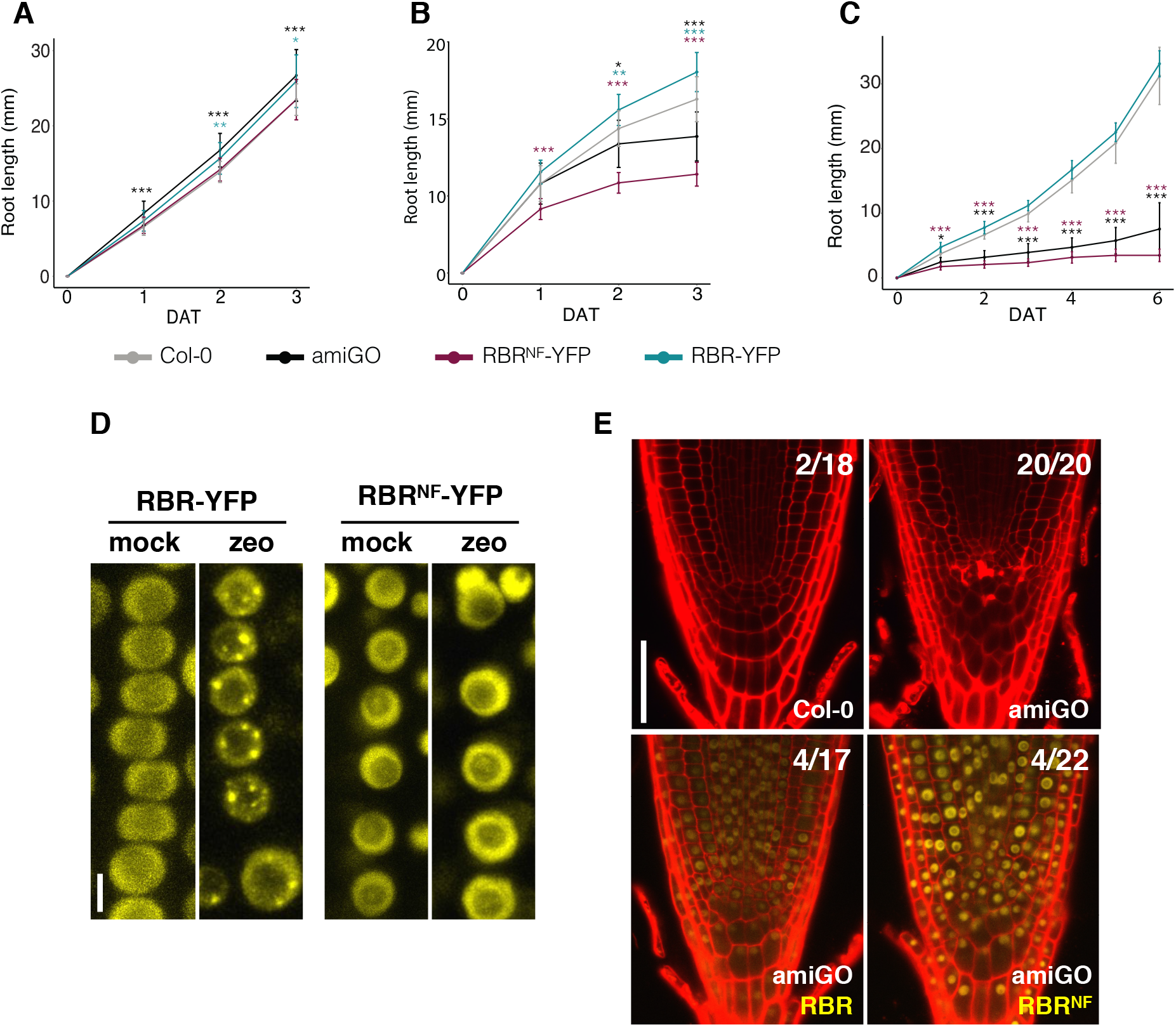
The LXCXE motif interacting domain of RBR is needed to cope with DNA damage. (A-C) Root growth comparison of Col-0, amiGO, RBR-YFP and RBRNF-YFP. Seedlings germinated and grown on 0.5 GM medium for 4-5 days were transferred to 0.5 GM medium without (A) or with (B) 3 μg/mL zeocin (zeo) for 3 days after transfer (dat), or incubated on 10μM zeo for 20 h and transferred again to 0.5 GM for recovery over 6 dat (C). Data in A,B) presented as mean +SD of two independent replicates, in C) a single replicate is presented. 10<n<17. ***p < 0.001, **p < 0.01, *p < 0.05. (D) Representative maximum-intensity projections of z-stack images from RBR-YFP and RBR^NF^-YFP living roots nuclei after 16 h incubation in 0.5 GM medium supplemented with without (mock) or with 10μg/mL zeo. (E) Cell death visualized by confocal imaging of longitudinal sections of propidium iodide (PI)-stained root tips 8 days post germination (dpg) on 0.5 GM medium without zeo; numbers indicate roots presenting dead cells in Col-0, amiGO and amiGO complemented with RBR-YFP or RBR^NF^-YFP. Scale bars, 5 μM in (D), 50 μM in (E). See also Figure S1.

Since RBR aggregates in nuclear foci with histone γH2AX and repair proteins RAD51 and BRCA1 upon induced DSB (Biedermann et al., 2017; Horvath et al., 2017), we asked whether the inability of RBR variants to recover from DNA damage relates to this process. Strikingly, focus formation was abolished in RBR^NF^-YFP (Fig 1D), suggesting that an LXCXE-containing protein is required to tether RBR to DSB. A conserved LXCXE motif in RAD54 (fig S1A), a DNA repair protein that also forms repair foci with RAD51 in Arabidopsis (Hirakawa and Matsunaga, 2019; Hirakawa et al., 2017), led us to test whether RAD54 might recruit RBR to foci. We failed to demonstrate that RBR and RAD54 bind directly (Fig. S1B), and no co-localization of RAD54 and RBR zeocin-induced foci was observed, in contrast to the highly co-localized RBR/E2FA foci (Fig S1C-H). Since RBR forms foci in the absence of RAD54 (Fig S1I), RAD54 does not recruit RBR to the DNA damage sites. In sum, the LXCXE-binding cleft is crucial for RBR foci formation and tethering by an as yet unidentified protein.

Root tips with reduced RBR levels display cell death even in unchallenging conditions, likely due to intrinsic genome instability (Biedermann et al., 2017; Cruz-Ramírez et al., 2013; Horvath et al., 2017; Wildwater et al., 2005), but whether this phenotype relates to defective foci dynamics is unknown. Since RBR^NF^-YFP rescued this phenotype observed in amigo roots to the same extent than RBR-YFP (Fig 1E), we conclude that the ability of RBR to form nuclear foci is dispensable to promote cell survival in standard growth conditions.

### Phosporylation state independent interaction of RBR with NAC044 through a conserved LXCXE motif

In a Y2H screening of the Arabidopsis transcription factors library (Pruneda-Paz et al., 2014) we identified NAC044 as a strong RBR interactor (preprint:(Zamora-Zaragoza et al., 2021)). NAC044 is a direct transcriptional target of SOG1 and, along with NAC085, the closest homolog of SOG1 (Ogita et al., 2018; Takahashi et al., 2019). We noticed that NAC044 contains an LXCXE motif in the C-terminus (Fig 2A,B); such motif is conserved among NAC044 orthologs in monocots and dicots species but is absent in NAC085 and modified in SOG1 (Fig 2A). Noteworthy, the NAC044 LXCXE motif is identical to that of RAD54 (compare Fig 2A and S1A). To test the RBR binding capacity of the NAC044 LXCXE motif, we performed Y2H assays. Figure 2C shows that E2FC binds to RBR and RBR^NF^, but NAC044 failed to interact with RBR^NF^. When the LXCXE motif in NAC044 was changed into GXCXG (hereafter NAC044^GCG^) the interaction with RBR was abolished (Fig 2C), confirming a similar experiment published recently (Lang et al., 2021). Split-luciferase assays showed that, while E2FA, E2FB and E2FC interacted *in planta* with both RBR and RBR^NF^ (Fig S2A), NAC044-RBR binding was suppressed by either RBR^NF^ or NAC044^GCG^ mutations (Fig 2D). Moreover, RBR and NAC044 interacted in a LXCXE-dependent manner upon zeocin treatment as shown by a split-lucieferase assay in stable transgenic Arabidopsis seedlings (Fig. S2B). In contrast, SOG1 was unable to interact with RBR (Fig 2C).

**Figure 2.**
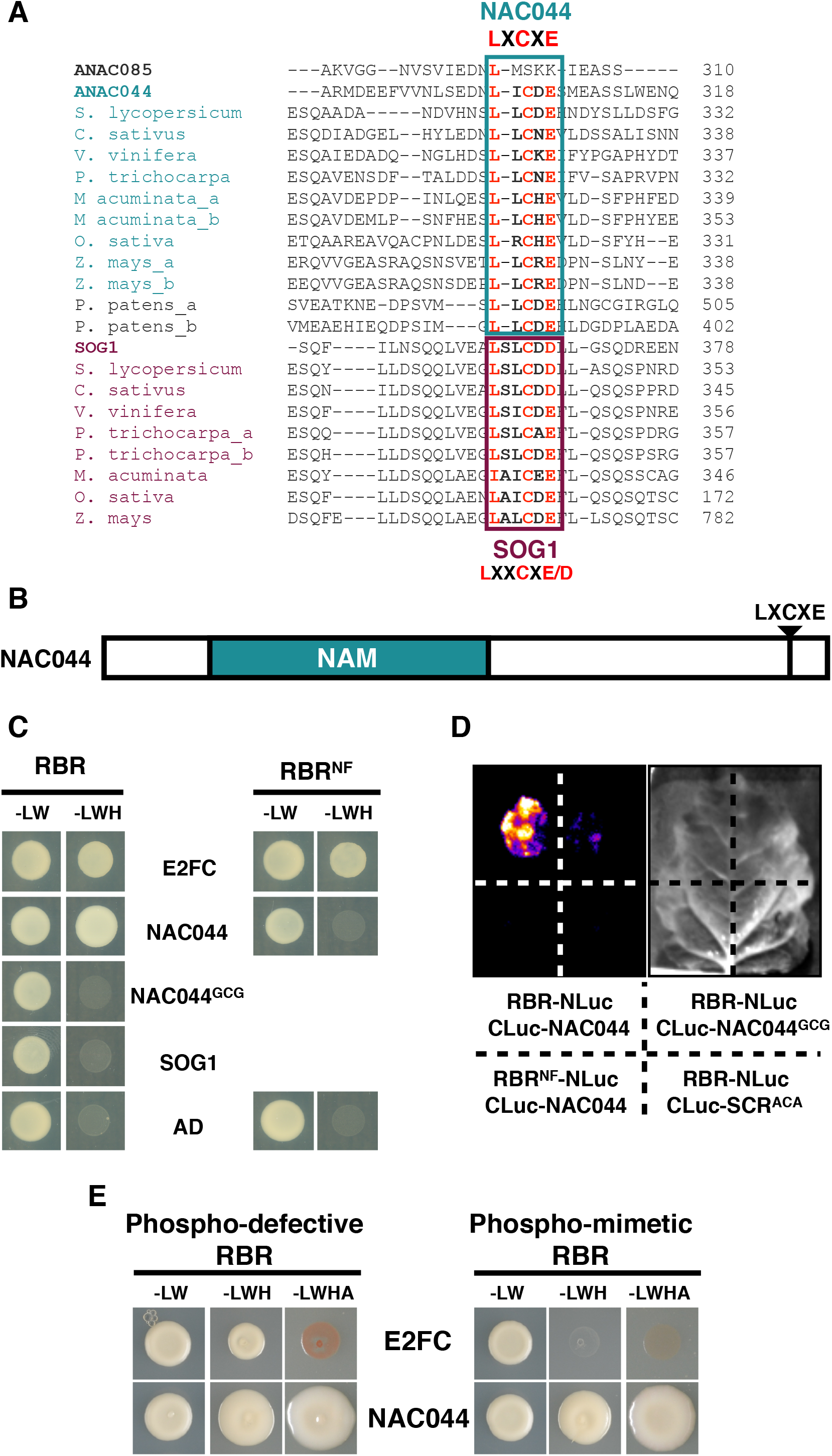
RBR interacts with NAC044 through a conserved LXCXE motif regardless RBR phosphorylation state. (A) Protein sequence alignment of Arabidopsis NAC044 (ANAC044) and SOG1 orthologs in the indicated plant species showing the fragment with LXCXE and LXXCXE/D motifs highlighted. No ANAC085 orthologs lacking an LXCXE motif were found in these species. (B) Schematic representation of NAC044 protein organization showing the relative positions of the NAM domain and LXCXE motif. (C) Yeast two-hybrid analysis showing that BD-RBR, but not BD-RBR^NF^, interacts with AD-NAC044, and neither of them binds AD-SOG1 nor an LXCXE-to-GXCXG-mutated AD-NAC044 protein (NAC044^GCG^). AD-E2FC and empty vector (AD) are positive and negative controls, respectively. Transformant yeast were dropped onto SD/-L/-W (-LW), SD/-L/-W/-H/+1.5mM3AT (-LWH). AD and BD, GAL4 activation domain and DNA binding domain, respectively. (D) Split Luciferase assay of RBR binding to NAC044 *in planta*. *N. benthamiana* leaves were coinfiltrated with the plasmid combinations and in the order indicated by dashed-line divided quadrants. Luciferase activity and a bright field images are shown. Representative images of 6 independent replicates. (E) Yeast two-hybrid analysis reveals that NAC044 interacts with a non-phosphorylatable RBR mutant (phospho-defective RBR) and phospho-mimmetic RBR. All 16 putative CDK-phospho-sites in RBR were mutated to Ala (phospho-defective) or to either Asp or Glu (Phospho-mimmetic). Transformant yeast were dropped onto SD/-L/-W (-LW), SD/-L/-W/-H/+1.5mM3AT (-LWH), SD/-L/-W/-H/-A (-LWHA). Data shown are representative of 3 independent replicates and belongs to a larger Y2H screening reported in (preprint: (Zamora-Zaragoza et al., 2021). See also Figure S2

While both E2FC and NAC044 interacted with a fully phospho-defective RBR, a phospho-mimetic RBR disrupted the binding to only E2FC (Fig 2E). Since NAC044 fosters stress-induced cell death and G2/M cell cycle arrest (Takahashi et al., 2019), and RBR is phosphorylated at the G1/S-phase transition (Boniotti and Gutierrez, 2001; Nakagami et al., 2002), there is a logical explanation for the ability of NAC044 to bind the hyper-phosphorylated form of RBR. Altogether, our results demonstrate that RBR interacts with NAC044 in a LXCXE-dependent manner but independent of RBR phosphorylation state –which is relevant in the likely scenario where both proteins act together after the G1/S phase transition.

### RBR is rapidly recruited to foci before NAC044 expression is induced by DNA damage

Is NAC044 the protein that recruits RBR to foci? we first generated transcriptional and translational reporters. We utilized a promoter that comprises the intergenic region and harbors the SOG1 binding site, a MSA element (mitosis-specific activator) and a E2F box (Fig S3A). pNAC044::GUS and pNAC044::3GFP-NLS resemble the previously reported expression pattern in the root tip (Takahashi et al., 2019), characteristic of cell cycle-regulated, and DNA damage-induced genes (Fig S3B) –contrasting with its age-dependent expression in the floral organ abscission zone (Fig. S3C).

pNAC044::NAC044:GFP (hereafter referred to as NAC044-GFP) never formed foci after a zeocin treatment that promoted RBR focus formation and NAC044-GFP accumulation in the same nuclei (Fig S3D), indicating that focal RBR is unable to bind NAC044. Moreover, RBR formed foci in the absence of NAC044 (Fig. S3E). Therefore, NAC044 is not the recruiter of RBR at foci, and more likely acts with RBR in the later transcriptional response rather than in the structural aspect of DNA repair. These roles are separated in the sub-nuclear space by LXCXE-binding constraints.

RBR repair functions are separated in time as well. RBR focus formation started immediately after a short exposure to zeocin and rapidly increased within 1h (Fig. 3A,B), whereas a long treatment induced a large amount of RBR foci that cleared out almost completely within 8 hours of recovery (Fig 3C,D). Noteworthy, histone γH2AX focus formation and clearance kinetics are similar (Charbonnel et al., 2011). Conversely, the pNAC044 reporters increased only after 3 hours following a short zeocin treatment, peaking after 12h, and lasting for at least 3 days (Fig S3B). These results indicate that RBR is recruited to foci well before NAC044 is highly induced, preventing their interaction during the early stages of the DDR, but released to the nucleoplasm where, later on, both proteins are present.

**Figure 3.**
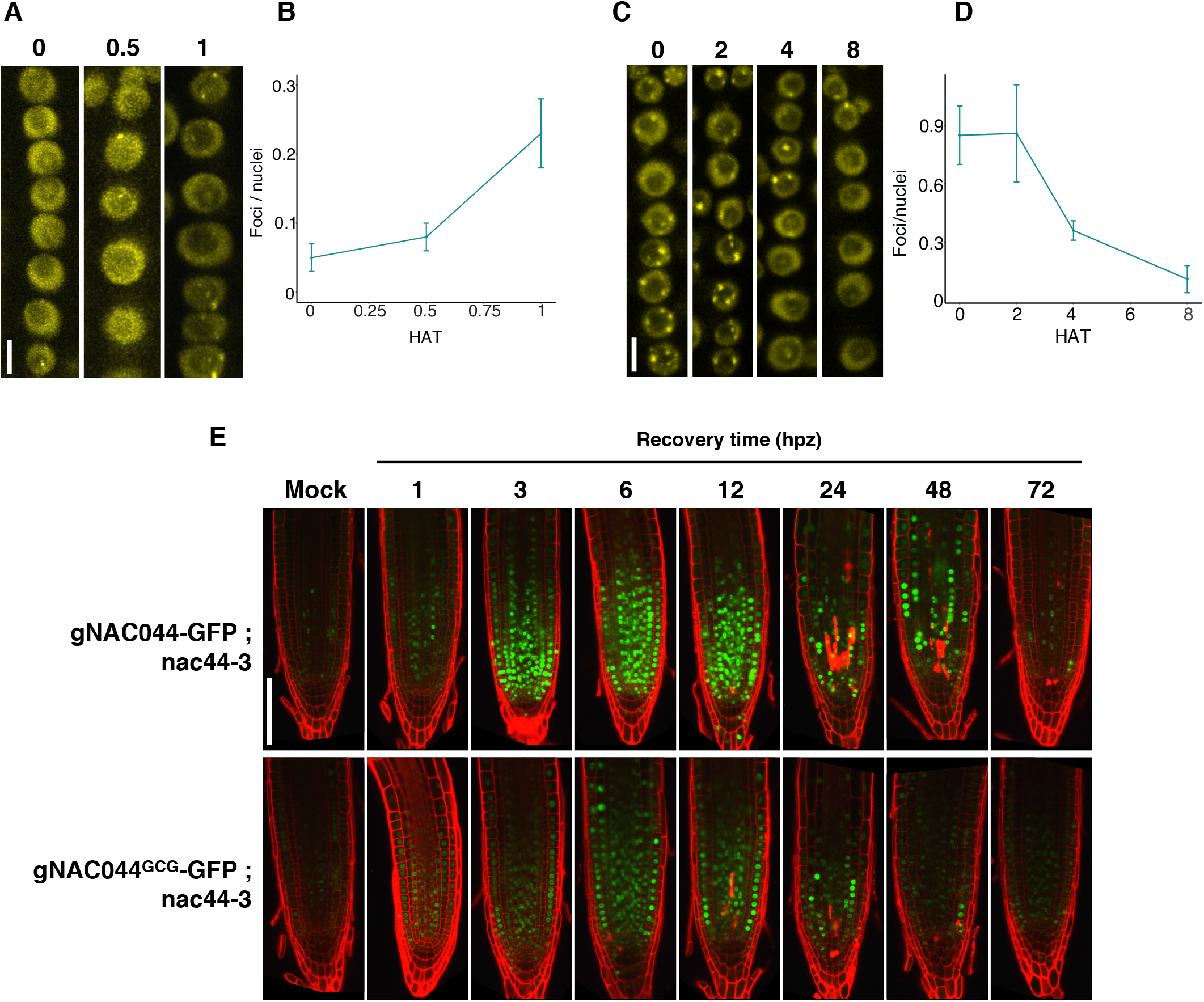
RBR is rapidly recruited to foci before NAC044 expression is induced by DNA damage. (A-D) RBR-YFP foci at 0, 0.5 and 1 h (A,B) or 0, 2, 4, 8 h (C,D) after zeo treatments: 20 μg/mL zeo for 2h (A,B) and 10 μg/mL zeo for 16h (C,D). Seedlings were transferred to 0.5 GM for the indicated recovery time (hours after zeo treatment, hat) before imaging. Representative maximum-intensity projections of z-stack images from RBR-vYFP living roots nuclei (A,C) and quantification of nuclear foci divided by the number of nuclei (B,D). Data in (B,D) presented as mean +SD. In (B) n > 3 roots, total nuclei per time point >1000; in (D) n>4, total nuclei per time point >2000. (E) Representative confocal images of longitudinal sections of roots from *nac044-3* complementing translational fusions pNAC044::gNAC044:GFP and pNAC044::gNAC044^GCG^:GFP after 2h incubation on 20 μg/mL zeo. Seedlings were transferred to 0.5 GM for the indicated recovery time before imaging. Scale bars, 5 μM in (A,C), 100 μM in (E). See also Figure S3.

### NAC044 interacts with RBR to induce cell death after DNA damage

To further explore the biological relevance of NAC044-RBR interaction, we generated both NAC044-GFP and its LXCXE-mutated version (pNAC044::NAC044^GCG^:GFP; hereafter NAC044^GCG^-GFP) in the genetic background of *nac044* knock-out alleles —*nac044-3* (Fig S3F) and the reported *nac044-1,* both of which exhibited a similar insensitivity phenotype to DNA damage (Takahashi et al., 2019)(Fig S3G,H). NAC044-GFP and NAC044^GCG^-GFP fully complemented *nac044-3* when seedlings grew on sustained zeocin conditions (Fig S3I). After a short zeocin exposure all reporters were induced, but the translational fusions displayed a broader expression domain than the promoter reporter (Fig 3 E; Fig S3B) —maybe due to different protein stability or transcriptional regulatory elements within the gene body. Both NAC044-GFP and NAC044^GCG^-GFP gradually accumulated, reaching maximum levels between 6-12h, then they gradually decreased over the course of two days (Fig 3 E). Along with NAC044 accumulation, the meristem shrinks and cell death increases during the first 24h.

We further tested the effect of a prolonged pulse of zeocin on root growth to address the physiological role of the RBR-NAC044 interaction. Besides the strong effect of RBR^NF^ in combination with reduced endogenous RBR levels showed in Figure 1C, the RBR^NF^–YFP prevented roots to resume growth, even in the presence of the endogenous RBR or in the absence of NAC044. Conversely, extra copies of Wt RBR seemed to have a positive effect on root growth (Fig S4A). Taken together, RBR^NF^ is not only less active due to its inability to bind LXCXE proteins, but has dominant features that prevent the root from coping with DNA damage. Therefore, failure to form foci and to interact with NAC044 can only partially explain RBR^NF^’s defect. As previously determined (Takahashi et al., 2019), *nac044-1* kept growing after zeocin treatment (Fig S4B), whereas several independent NAC044-GFP and NAC044^GCG^-GFP transgenic lines with varying accumulation levels (Fig S4C) consistently showed over-complementation of *nac044* mutants, suggesting that the function of NAC044 on root growth is influenced by the GFP fusion. Moreover, NAC044^GCG^-GFP behaved similar to NAC044-GFP, indicating that the interaction with RBR does not influence the suppressive function of NAC044 on root growth as proposed recently (Lang et al., 2021).

NAC044 has a second function, as it also mediates SOG1-dependent cell death upon DNA damage (Takahashi et al., 2019). As cell death in standard and damaging conditions increases when RBR levels are down regulated, we asked whether NAC044 and RBR functions might converge in this process. After 24 h of treatment with zeocin, the cell death area was smaller in the *nac044-1* mutant and larger in the amiGO root tips as compared to Wt (Fig 4). Noteworthy, NAC044-GFP but not NAC044^GCG^-GFP completely restored the induction of cell death in *nac044-1.* Furthermore, the introgression of *nac044-1* in the amiGO line significantly reduced the cell death (Fig 4). Altogether, our results indicate that NAC044 binds to RBR to inhibit the latter from promotion of cell survival upon DNA damage.

**Figure 4.**
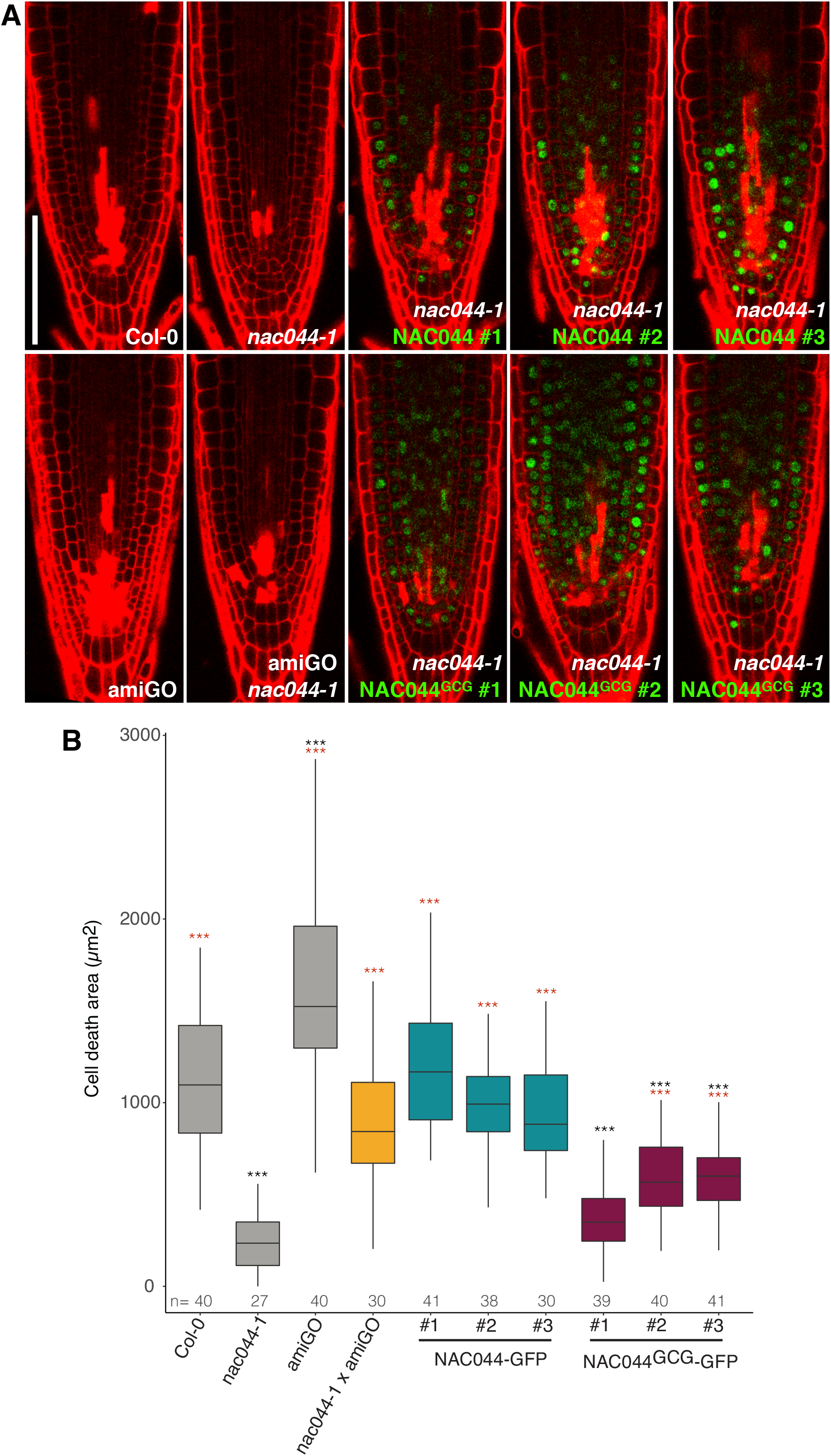
NAC044 interacts with RBR to induce cell death after DNA damage. (A) Cell death visualized by PI accumulation in a representative confocal image of longitudinal sections of root tips from the indicated genotypes. Three independent transformant lines of NAC044-GFP and NAC044^GCG^-GFP in *nac044-1* mutant background were tested. 4 dpg seedlings were incubated on 10 μg/mL zeo for 24h before imaging. (B) Box plots of quantified cell death area related in the root tips mentioned in A). Data presented as median and interquartile range from two biological replicates, n denotes total number of scored roots. A third biological replicate showed similar results. Wilcoxon test, black and red asterisks denote ***p < 0.01 as compared to Col-0 or *nac044-1,* respectively. Note that all NAC044^GCG^ but no NAC044-GFP lines are different than Col-0, and one NAC044^GCG^ line is not different than *nac044-1*. Scale bar, 100 μM. See also Figures S6 – S8.

## Discussion

We report here that the coordinated activities of RBR during the DDR are separated in space and time, establishing an orderly series of events that couples cell survival with DNA integrity. Quickly after DNA injury, RBR is recruited to nuclear foci by a protein-protein interaction requiring the B-pocket subdomain and, most likely, an LXCXE-containing protein. Meanwhile, the induction of the LXCXE-containing protein NAC044 arrests cell proliferation and growth. If the damage persists, NAC044 represses the cell protective activity of RBR to promote cell death. We postulate that the dynamic aggregation and release of RBR from nuclear foci enables a spatio-temporal separation of events that determines the decision of resuming growth or undergoing cell death.

Under unchallenging growth conditions RBR-^NF^ completely restored the compromised cell survival after down regulating endogenous RBR. In animals, pRb is recruited to nuclear foci in a E2F1-dependent manner and pRb foci kinetics are similar to those of histone γH2AX (Vélez-Cruz et al., 2016). Strikingly, RBR^NF^-YFP was unable to accumulate in nuclear foci despite its ability to interact with E2F proteins, suggesting that in plants, a LXCXE-containing protein recruits RBR to the DNA damage site. Accordingly, the *rbr1-2* allele, which lacks the B-pocket sub-domain, displays reduced sister chromatide cross-over and DSB-dependent foci formation of meiotic proteins during prophase I (Chen et al., 2011), highlighting the importance of the B-pocket-mediated interactions in programmed and stress-induced chromatin repair. We conclude that, when favorable growth conditions are maintained, the ability to interact with LXCXE proteins is somehow dispensable, but when DNA damage occurs, LXCXE-mediated interactions become essential.

DNA repair foci are highly dynamic (Biedermann et al., 2017; Gentric et al., 2020; Nakamura et al., 2010; Polo and Jackson, 2011). Once at damage sites, RBR recruits and/or co-localizes with other proteins like E2F, F-BOX-LIKE PROTEIN 17 (FBL17) and the repair proteins BRCA1 and RAD51 (Biedermann et al., 2017; Gentric et al., 2020; Horvath et al., 2017). RBR nuclear foci may display a different composition and dynamics according to the cell cycle phase at the time when damage is perceived. Further research is required to discover the protein that recruits RBR to foci, and to characterize dynamic RBR focus composition. The structural role of RBR could be further elucidated using a null *rbr* background for RBR^NF^-YFP to follow the DNA repair process.

Our data reveal that RBR aggregates in nuclear foci during the first hour after DNA damage induction, well before NAC044 accumulates in root meristem cells. Nearly 90% of the RBR foci are cleared during the first 8 h of recovery, simultaneously with the NAC044 accumulation peak. Whether focus-released RBR immediately binds NAC044 is not clear, but the effects of disrupting the LXCXE motif in NAC044 are already evident during the first 24 hours after DNA damage, when cell death becomes visible. Apparently, the main biochemical constraint for NAC044 to bind RBR is the availability of the B-pocket sub-domain, while the phosphorylation state of RBR seems irrelevant for binding. However, the exact timing of the RBR-NAC044 interaction remains to be determined. Moreover, RBR-NAC044 dimers probably establish other protein interactions that influence the transcriptional outcome of the complex. On the one hand, NAC044 is expected to interact with its closest homologs NAC085 and SOG1, as the single, double and triple mutants have similar effects on cell death (Takahashi et al., 2019); and NAC044 also binds to the DREAM complex component LIN37 upon DNA damage (Lang et al., 2021). On the other hand, the dominant effect of RBR^NF^-YFP likely reflects other non-LXCXE interactions of RBR, for example with E2FA. Whether these interactions occur simultaneously to NAC044 binding remains an open question.

NAC044 controls cell division by promoting accumulation of repressor MYB3R proteins in response to DNA damage (Takahashi et al., 2019), which in turn bind to the MSA element of G2/M-specific gene promoters to arrest cell cycle progression (Bourbousse et al., 2018; Chen et al., 2017; Ito et al., 1998, 2001; Kobayashi et al., 2015). The mechanism by which NAC044 promotes cell death is less clear, but also includes the action of repressor MYB3R proteins (Chen et al., 2017; Takahashi et al., 2019). Repressor MYB3R proteins are components of the Arabidopsis DREAM complex, a multimeric protein complex containing among others RBR and E2F proteins (Kobayashi et al., 2015; Ning et al., 2020). Thus, the SOG1 and RBR DDR pathways may use this protein complex to couple cell cycle and cell death decisions to DNA integrity. A recent publication shows the dynamic composition of the DREAM complex upon DNA damage induction, and reveals that mutants in the DREAM complex members E2FB and LIN37 exhibit defective root growth arrest, similar to nac044 mutants (Lang et al., 2021). While we cannot exclude the possibility that NAC044 cooperates with the DREAM complex to restrain root growth, our results indicate that the direct NAC044-RBR interaction is more relevant to cell survival.

The *NAC044* gene displays a DNA damage and cell cycle-dependent expression pattern, MSA, E2FA and SOG1 binding elements reside in its promoter, and *NAC044* transcription is regulated by SOG1, MYB3R3, E2FA and RBR (Bourbousse et al., 2018; Bouyer et al., 2018; Kobayashi et al., 2015; Ogita et al., 2018; Verkest et al., 2014). These observations raise the question whether different RBR protein complexes regulate DDR promoters independently, or converge in larger transcriptional regulatory complexes than previously thought. In animals, the SOG1 functional analog, p53, cross talks with the RBR and DREAM complex pathways during DDR. In this process, p53 induces the CDK inhibitor (CKI) p21 to facilitate the DREAM complex formation through hypo-phosphorylation of pRb-like proteins p130 and p107, promoting a transcriptional switch to repress DREAM complex targets (Engeland, 2018). Since SOG1 induces CKIs (Bourbousse et al., 2018; Ogita et al., 2018; Yi et al., 2014), it is possible that RBR plays a phosphorylation-regulated role in the DDR beyond the NAC044 interaction.

Altogether, our data indicate that RBR interaction with the LXCXE motif is not only essential for its structural and transcriptional functions during the DDR, but also imposes a timing mechanism to coordinate them in different sub-nuclear locations. Soon after damage is perceived, RBR is recruited to foci to aid DNA repair; the parallel activation of NAC044 triggers cell cycle arrest and cell death later on. The short-time dynamic accumulation of RBR in foci might create a flow of RBR between the damage site and the nucleoplasm where it can interact with NAC044. Both processes depend on the binding to LXCXE motif. The RBR-NAC044 interaction promotes cell death possibly by an inhibition of RBR protective function which, in turn seems to depend on binding E2FA (Horvath et al., 2017; Wang et al., 2014). Thus, LXCXE-mediated interactions of RBR lie at the interface between the DNA repair process and the life or death cellular decisions afterwards.

## Materials and Methods

### Plant material and growth conditions and treatments

*Arabidopsis thaliana* ecotype Col-0 was used as wild-type control. Mutant lines *nac044-1* (SAIL_1286_D02) (Takahashi et al., 2019), nac044-3 (WiscDsLox293-296invF22), and *rad54* (SALK124992) were obtained from The Nottingham Arabidopsis Stock Centre (NASC). RAD54-YFP (Hirakawa et al., 2017), E2FA-GFP (Magyar et al., 2012), amiGO-RBR (amiGO), pRBR::RBR:YFP (Cruz-Ramírez et al., 2013), and pRBR::RBR:mRFP (Cruz-Ramírez et al., 2012) lines were previously published. pRBR::RBR^NF^:YFP line was kindly donated by Sara Diaz-Trivino. Unless otherwise noticed, amiGO was used as background for RBR and RBR^NF^ transgenic plants, and *nac044-3* or *nac044-1* for NAC044:GFP and NAC044GCG:GFP as indicated. Homozygous pNAC044-GUS and pNAC044-3xGFP-NLS are in Col-0. Seeds were fume-sterilized in a sealed container with 100 ml bleach and 3 ml of 37% hydrochloric acid for 3–5 h; then suspended in 0.1% agarose, stratified for 2 d at 4 °C in darkness, plated on 0.5x Murashige and Skoog (MS) plus vitamins, 1% sucrose, 0.5g/L 2-(N-morpholino) ethanesulfonic acid (MES) at pH 5.8, and 0.8% plant agar, and grown vertically for the 4-6 d (as indicated in each figure) at 22°C with a 16h light/8h dark cycle. For DNA damaging treatments, a filter-sterilized 20 mg/mL stock solution prepared from zeocin powder (Duchefa Biochemie) was diluted to 3 μg/mL, 10 μg/mL, or 20 μg/mL in 0.5x MS + vits, 1% sucrose, 0.5g/L MES at pH 5.8, 0.8% plant agar before pouring into plates. Seedlings were transferred to zeocin-containing plates for 2, 16, or 24 h as indicated in figure legends before imaging or transfering back to fresh 0.5 GM medium.

### Plasmid construction and plant transformation

The constructs pNAC044::GUS, pNAC044::3xGFP:NLS, pNAC044::NAC044:GFP and pNAC044::NAC044GCG:GFP were cloned using the GreenGate system (Lampropoulos et al., 2013). Briefly, a 980 bp fragment upstream of the ATG was amplified with the primers pANAC044 F and pANAC044 R1 (translational fusion) or pANAC044 R2 (transcriptional fusions); for translational fusions, the genomic segment of NAC044 without stop codon (gNAC044) was amplified with the primer pair gANAC044 F / gANAC044 R. Mutagenesis of the of the LXCXE into GXCXG motif was done by amplifying gNAC044 in two fragments with primers pairs gANAC044 F/mut gANAC044 R and mut gANAC044 F/gANAC044 R. Promoter and gNAC044 PCR products were cloned together with the plasmids pGGD001, pGGE009 and pGGF001 (for translational fusion); with pGGB002, pGGC025, pGGD006, pGGE009, and pGGF001 (for 3xGFP-NLS transcriptional fusion); and with pGGB002, pGGC051, pGGD002, pGGE009, pGGF001 (for GUS transcriptional fusion) into the vector pGGZ003 by combining equimolar amounts of each part in 15 μL dig-lig reactions including 10mM ATP, 1x Green Buffer (Thermo Fisher), 1μL T4 DNA ligase (30u/μL), 1μL BsaI restriction enzyme (Thermo Fisher) according to the GreenGate protocol. Constructs were transformed into Arabidopsis plants by the flower dip method.

Previously reported entry and expression clones are listed in Table S2. The pEXP22-NAC044 expression clone, and the product of a two fragments-overlapping PCR with primers pairs AttB cNAC44 F/mut cNAC44 R and mut cNAC44 F/AttB cNAC44 R that introduce attB sites and mutagenize the LXCXE motif to GXCXG on NAC044 CDS, were recombined into pDONR221 vector with Gateway BP clonase II enzyme mix (Invitrogen) to generate the corresponding entry clones. The CDS of E2FA, E2FB and E2FC, amplified with the corresponding primers listed in Table S1. were cloned into pGEMt-easy 221 vector by BP clonase reaction. We generated pDEST-NLuc and pDEST-CLuc Gateway-compatible destination vectors by conventional cloning of the attR-flanked Gateway cassette amplified with the primers GWcass_SplitLUC_F and GWcass_CLuc_R or GWcass_NLuc_R (Table S1) and digested with BpiI restriction enzyme (Thermo Fisher) to generate sticky ends compatible with BamHI/SalI digested pCAMBIA-NLUC and pCAMBIA-CLuc vectors (Chen et al., 2008). Entry clones were recombined into pDEST22 or pDEST32 destination vectors for Y2H assays, and/or into pDEST-NLuc or pDEST-CLuc for Split-Luciferase assays with Gateway LR clonase II enzyme mix (Invitrogen).

### Microscopy and image processing

A 10 μg/mL Propidium Iodide (PI) staining solution (or mQ water for nuclear foci imaging) was used for whole-mount visualization of live roots with CLSM using a Zeiss LSM 710 system as described in (Zhou et al., 2019); PI, GFP, and YFP were visualized using wavelengths of 600-640 nm, 500-540, and 525-565 nm, respectively. Tissue clearing with ClearSee reagent was performed as described in (Kurihara et al., 2015). In brief, tissues were fixed with 4% w/v paraformaldehyde in 1x PBS for 30 min under vaccum (~690 mmHg) at room temperature, washed twice and submerged in the Clearsee solution (10% w/v Xylitol powder, 15% w/v sodium deoxycholate, and 25% w/v urea dissolved in water) for 1-2 week at room temperature. Images were taken with ZEN 2012 software (Zeiss) and processed with ImageJ. Brightness and contrast of the final figures was enhanced to the exact same values.

### Protein-proten interaction assays

Protein-Protein interactions by Y2H was performed using the ProQuest system (Thermo Scientific). Co-transformation of pEXP22 and pEXP32 expression clones into the PJ69-4A yeast strain was performed as described in (De Folter and Immink, 2011). Split-Luciferase assays in Nicothiana benthamiana leaves were performed as described in (Chen et al., 2008) using an exposure time of 7 min. For Split-LUC in Arabidopsis stable transformants, we used an exposure time of 10 min. Luciferase activity was detected after spraying with 1 mM D-luciferin (Duchefa Biochemie) using an (−80°C) air-cooled CCD Pixis 1024B camera system (Princeton Instruments, Massachusetts, USA) equipped with a 35mm, 1:1.4 Nikkon SLR camera lens (Nikon, Tokyo, Japan) fitted with a DT Green filter ring (Image Optics Components Ltd, Orsay, France) to block chlorophyll fluorescence. Lufiferase images were processed using a “Fire” lookup table in ImageJ, adjusting brightness and contrast to the same values in all images.

### Protein sequence retrieval and alignments

Arabidopsis thaliana NAC044, NAC085, and RAD54 protein sequences were retrieved from The Arabidopsis Information Resource (TAIR, arabidopsis.org) and used as query to BLAST search on Gramene.org for plant orthologs, or on NCBI for non-plant orthologs. Annotated orthologs or the top score hit sequences were retrieved and aligned with Clustal-omega (https://www.ebi.ac.uk/Tools/msa/clustalo/) to the Arabidopsis sequences.

### Quantification and Statistical analysis

Statistical parameters and tests are mentioned in figure legends. Calculations were done using GraphPad Prism software 5.0 (GraphPad Software, San Diego, Calif) and R-based statistical analyses.

## Acknowledgements

We thank to Prof. Lieven De Veylder (VIB-UGent Center for Plant Systems Biology) for his valuable suggestions on this manuscript, and for sharing plasmid material; to Prof. Sachihiro Matsunaga (Tokyo University of Science) for sharing the RAD54-EYFP line; Dr. Yessica Alina Rodríguez-Rosales (Radboud UMC) for her support with statistical analysis and R plots. WZ was supported by the National Natural Science Foundation of China (32070874); JZZ was supported by Consejo Nacional de Ciencia y Tecnología (CONACyT, Mexico, 383871).

## SUPPLEMENTARY FIGURES

### Supplementary figure legends

**Figure S1.**
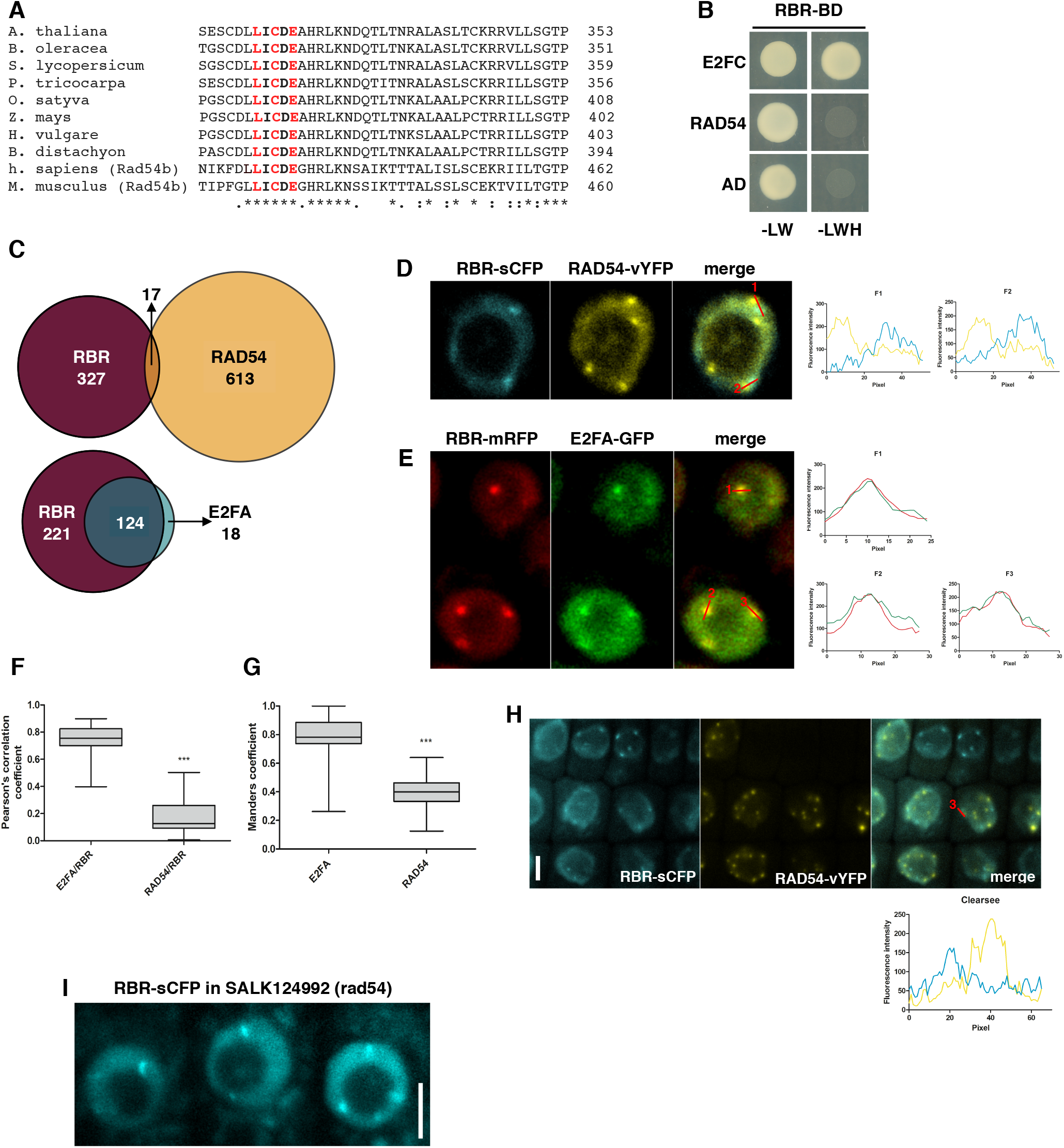
RAD54 and RBR act independently in DNA damage repair. (A) Protein sequence alignment of RAD54 orthologs in the indicated species showing the conserved LXCXE motif in red. (B) Yeast two-hybrid analysis showing that RAD54 and RBR do not interact. E2FC and empty vector (AD) are positive and negative controls, respectively. Transformant yeast were dropped onto SD/-L/-W (-LW), SD/-L/-W/-H/+1.5mM3AT (-LWH). AD and BD, GAL4 activation domain and DNA binding domain, respectively. (C) Venn diagrams showing the number of RBR foci overlapping with RAD54 or E2FA foci. Confocal images from >5 roots from the RBR-sCFP/RAD54-vYFP and RBR-mRFP/E2FA-GFP genotypes were used for pooled-counting after 16h incubation on 10 μg/mL zeo. The 17 RBR-sCFP/RAD54-vYFP overlapping foci resemble those highlighted in D), with only partial or no actual colocalization. (D,E) Confocal images (single section) of root cells showing colocalization of RAD54-vYFP and RBR-sCFP D) and E2FA-GFP and RBR-mRFP E). (F,G) Comparison of Pearson correlation coefficients of E2FA-GFP/RBR-mRFP vs RAD54-vYFP/RBR-sCFP (F) and Manders coefficients indicating the fraction of RBR-mRFP overlapping with E2FA-GFP vs the fraction of RBR-sCFP overlapping with RAD54-vYFP (G). Data presented as median and interquartile range, n=30 foci from >5 roots. (H) Maximum-intensity projection of z-stack confocal images of a Clearsee treated root showing RAD54-vYFP and RBR-sCFP foci after 16h incubation on 10 μg/mL zeo. (I) Confocal image (single section) of living root cells showing RBR-sCFP foci in rad54 T-DNA insertion allele SALK_124992 genetic background after 16h incubation on 10 μg/mL zeo. Representative image of >5 independent replicates. Fluorescence intensity profiles of the indicated foci in (D,E,H) show colocalization of RBR (red) and E2FA (green), or RBR (blue) and RAD54 (yellow).

**Figure S2.**
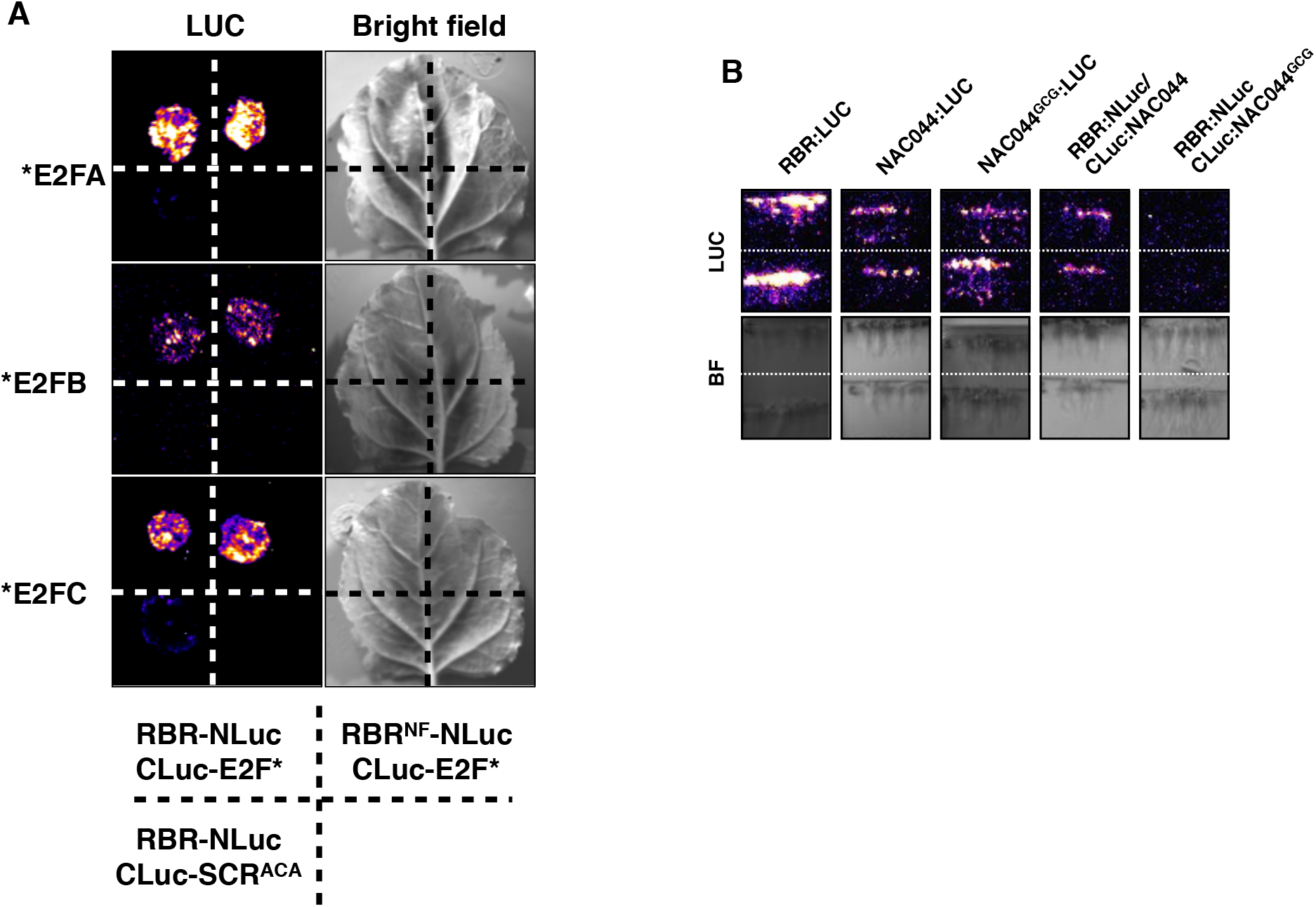
*In planta* Split-LUCIFERASE assays show that RBR^NF^ interacts with E2F proteins and NAC044 interacts with RBR through the LXCXE motif. (A) Split Luciferase assay showing that RBR^NF^ interacts with E2FA, E2FB and E2FC in *N. benthamiana* leaves that were co-infiltrated with the combinations and in the order indicated in quadrants. The LXCXE-to-AXCXA-mutated SCARECROW (SCR^ACA^) is used as negative control. LUC activity and the corresponding bright field images of a representative leaf from 3 independent replicates are shown. (B) Luciferase activity and the corresponding bright field images of two independent Arabidopsis T2 stable transformant lines (divided by a dotted line) expressing the constructs indicated in the schematic representation at the bottom. Since the constructs contain a FAST-R red-seed coat selection marker, only seeds containing the constructs were germinated on 0.5GM medium plates. 4 dpg, seedlings were treated with 0.5 GM liquid medium supplemented with 20 μg/mL for 2h and all lines were imaged together 24h later. Note that the three first images from the left contain single gene fusions with a full length LUCIFERASE tag, while Split-LUC constructs between RBR and NAC044 or NAC044^GCG^, are showed in the last two images. The number of independent transformant lines for each construct was 6 to 8, and 10 for RBR-NLuc/CLuc-NAC044^GCG^, all displaying similar results.

**Figure S3.**
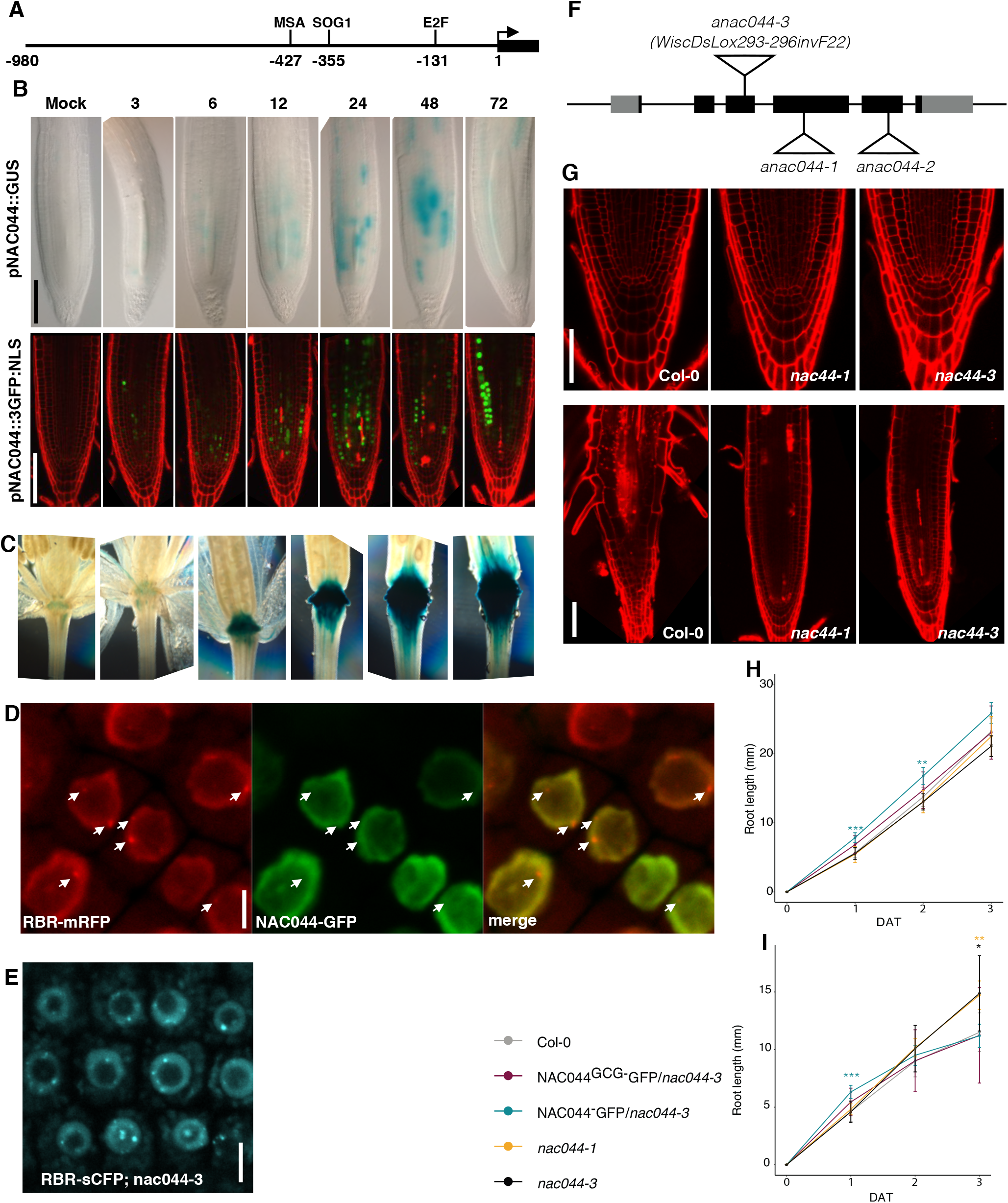
NAC044 is expressed after DNA damage, does not recruit RBR to nuclear foci and inhibits root growth. (A) Schematic representation of the NAC044 promoter cloned for transcriptional and translational fusions that was sufficient to recapitulate the previously reported expression pattern of NAC044 and to complement the *nac044* null mutants. The position of the MSA, SOG1 and E2F elements are indicated. (B) NAC044 promoter activity at the indicated times after mock treatment or 2h incubation on 20 μg/mL zeo as shown by transcriptional fusions with GUS (pNAC044::GUS) and a nuclear localized 3xGFP (pNAC044::3GFP:NLS). (C) Age dependent pNAC044::GUS activity in the abscission zone of inflorescences. (D) Confocal image (single section) of a Clearsee treated root showing NAC044-GFP and RBR-mRFP colocalization in nuclei after 16h incubation on 10 μg/mL zeo. White arrows indicate the position of RBR foci in nuclei with NAC044-GFP signal. Representative image of >5 independent replicates. We never observed NAC044-GFP foci. (E) Confocal image (single section) of living root cells showing RBR-sCFP foci in *nac044-3* genetic background after 16h incubation on 10 μg/mL zeo. Representative image of >5 independent replicates. (F) Schematic representation of the NAC044 gene showing the T-DNA insertion sites of *nac044-1*, *nac044-2* and *nac044-3*. Exons and introns are represented as black boxes and lines, respectively. (G) Confocal images of longitudinal sections of PI-stained root tips from 4-5 dpg Col-0, *nac044-1* and *nac044-3* seedlings germinated and grown in 0.5 GM medium (top panel), or transferred to 0.5 GM supplemented with 3 μg/mL zeo for 3 days (lower panel). (H,I) Root growth comparison of Col-0, *nac044-1,* and *nac044-3* complemented with pNAC044::NAC044:GFP (NAC044-GFP) and pNAC044::NAC044^GCG^:GFP (NAC044^GCG^-GFP). Seedlings germinated and grown on 0.5 GM medium for 5 days were transferred to 0.5 GM medium without (H) or with (I) 3 μg/mL zeo for 3 days. Data presented as mean +SD of a single replicate (due to high variability between independent replicates), 14>n>10. ***p < 0.001, **p < 0.01, *p < 0.05 (Student’s t-test). Scale bars, 100 μM in (B,C), 5 μM in (D,E), 50 μM (G).

**Figure S4.**
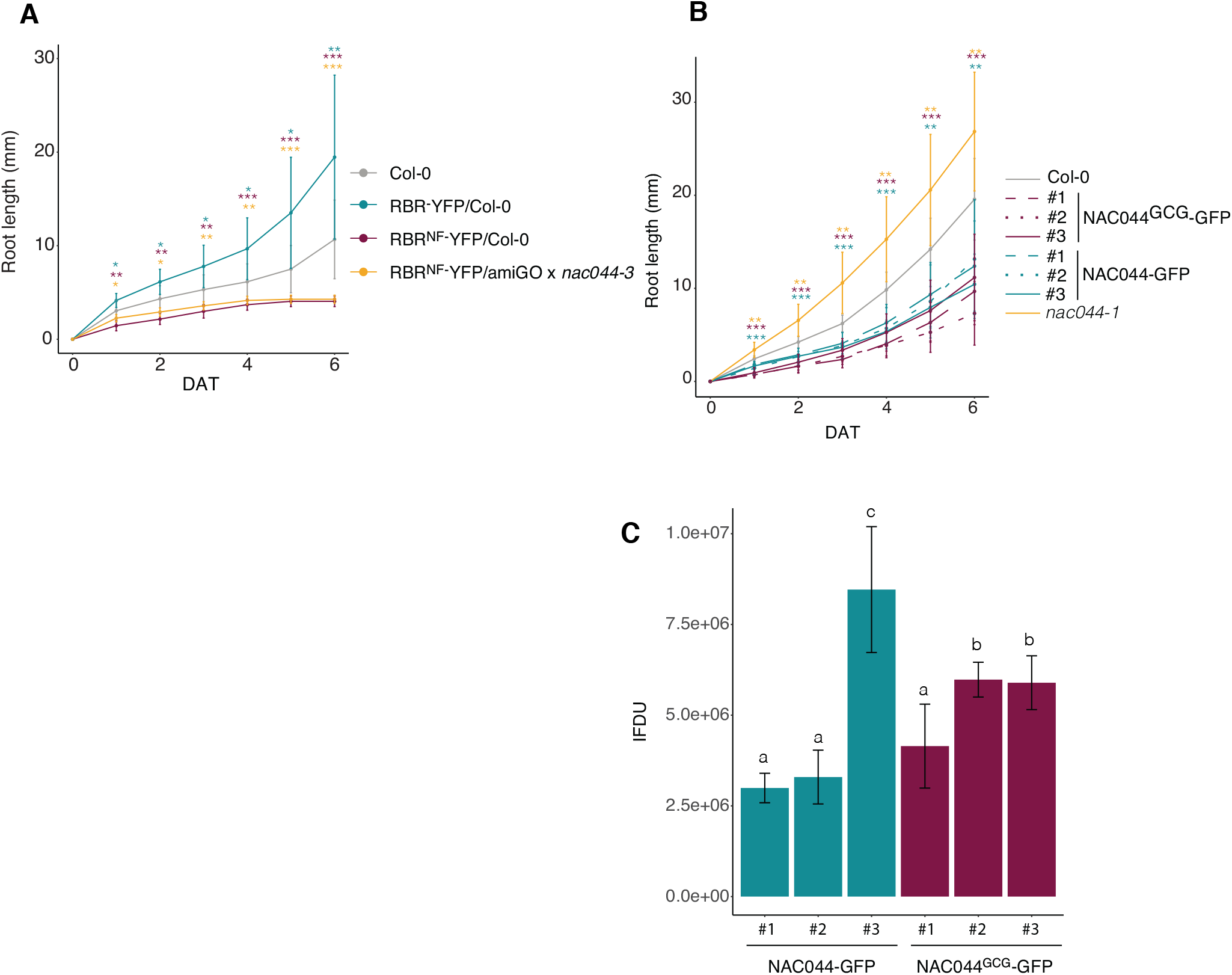
RBR^NF^ dominantly affects the DNA damage response through a different mechanism than binding NAC044. (A,B) Root growth after DNA damage. 4 dpg seedlings were incubated on 10 μg/mL zeo for 20-24 h and transferred again to 0.5 GM medium plates for recovery over 6 days. Due to the variability between replicates, a single representative biological replicate is shown. Root growth was scored for 6 days after zeo treatment (dat). Genotypes in (A) are Col-0, homozygous RBR-YFP in Col-0 background, homozygous RBR^NF^-YFP in Col-0, and homozygous F3 generation of RBR^NF^-YFP in amiGO background crossed with *nac044-3;* in (B) Col-0, *nac044-1,* three independent transformant lines of NAC044-GFP in *nac044-1,* and three independent transformant lines of NAC044^GCG^-GFP transformed in *nac044-1 (similar results obtained in the nac044-2 background).* In B) the complementing NAC044-GFP and NAC044^GCG^-GFP form a single statistically similar group. Data presented as mean +SD of a single replicate (due to high variability between independent replicates); n>12. ***p < 0.001, **p < 0.01, *p < 0.05 (Wilcoxon signed rank test). (C) Bar graph showing the quantification of integrated fluorescence density of the transformant lines of NAC044-GFP and NAC044^GCG^-GFP in *nac044-1* mentioned in (B) and Figure 4. 4 dpg seedlings were incubated on 10 μg/mL zeo for 24h before imaging. At least 8 roots per line were used for quantification using imageJ software. Lowercase letters denote ststistically different groups p<0.05 (Kruskal-Wallis).

